# Antimicrobial susceptibility testing for diverse *Malassezia* species

**DOI:** 10.1101/2022.12.09.519789

**Authors:** Brooke Rathie, Bart Theelen, Martin Laurence, Rebecca S. Shapiro

**Affiliations:** Department of Molecular and Cellular Biology, University of Guelph, Guelph, Ontario, Canada N1H 5N4; Westerdijk Fungal Biodiversity Institute, Utrecht, The Netherlands; Malassezia Foundation, Montreal, Quebec, Canada

**Author notes:** Correspondence &.

## Abstract

The genus *Malassezia* is an opportunistic lipid-dependent yeast that is associated with common skin diseases and has recently been associated with Crohn’s disease and certain cancers. Understanding the susceptibility of *Malassezia* to diverse antimicrobial agents is crucial for identifying effective antifungal therapies. Here, we tested the efficacy of isavuconazole, itraconazole, terbinafine and artemisinin against three *Malassezia* species: *M. restricta, M. slooffiae*, and *M. sympodialis*, using microbroth dilution techniques. We found antifungal properties for the two previously-unstudied antimicrobials: isavuconazole and artemisinin. Overall all *Malassezia* species were particularly susceptible to itraconazole, with a MIC range from 0.0015 to 0.1562 μM.

**Importance:** The *Malassezia* genus is known to be a cause of skin conditions and has recently been associated with diseases such as Crohn’s disease, pancreatic ductal carcinoma and breast cancer. This work was completed to assess the susceptibility to a variety of antimicrobial drugs on diverse *Malassezia* species, in particular *Malassezia restricta*.

## Introduction

The most prevalent fungal genus in the human skin microbiome is *Malassezia* (1). These fungi are known agents in human skin disorders such as dandruff and seborrheic dermatitis, as well as systemic infections in immunocompromised individuals (2, 3). It has been convincingly shown that *Malassezia*, especially *M. restricta*, are highly prevalent in human mucosal surfaces such as the mouth, gut, nose, pancreas and vagina *(4–6)*. Recent studies have implicated *Malassezia* in Crohn’s disease (7), pancreatic ductal carcinoma (8) and breast cancer (9), raising the possibility that reducing or eliminating *Malassezia* from affected organs might improve clinical outcomes of these diseases. The aim of this study was to evaluate the *in vitro* activity of diverse antimicrobial agents against three *Malassezia* species: *M. restricta, M. sympodialis* and *M. slooffiae*. To our knowledge, this is the first study to examine the effects of isavuconazole and artemisinin on the *Malassezia* genus.

We studied the susceptibility of four antimicrobials: itraconazole, isavuconazole, terbinafine and artemisinin. Itraconazole and isavuconazole are both triazoles which inhibit lanosterol 14α-demethylase, an enzyme necessary for the biosynthesis of ergosterol, the critical sterol of the fungal cell membrane. Terbinafine is an allylamine that inhibits squalene epoxidase, an enzyme which catalyzes the conversion of squalene to lanosterol in the ergosterol synthesis pathway. Artemisinin is an antimalarial drug, which has also been reported to have fungistatic activity (10–12). The mechanisms of action of artemisinin against fungi is still unknown. However, there has been research showing artemisinin influences Ca^2+^ ATPases in *Plasmodium falciparum, Saccharomyces cerevisiae* and *Candida glabrata* (12).

Antifungal susceptibility testing of *Malassezia* has been limited by the inability to reliably utilize conventional yeast microbiology protocols (13). Comparisons between minimum inhibitory concentration (MIC) experimental assays have proven to lack congruence (14, 15). Past studies have exploited large strain numbers to determine a general trend for *Malassezia* susceptibility to antifungals, including itraconazole amongst a variety of azoles, as well as terbinafine and amphotericin B (14–17). While there are multiple studies which have examined faster-growing *Malassezia* species MICs, few have examined the slow-growing *Malassezia* species such as *M. restricta* (15, 17).

Turbidity readings of *Malassezia* can underestimate growth and are unreliable due to the fungi’s tendency to clump. Past studies have used a colorimetric MIC method to overcome this issue for the *Malassezia* genus, utilizing resazurin to measure cell metabolism during or after antifungal treatment (15, 18, 19). Here, we built upon techniques used in these past studies to develop an effective process to measure *Malassezia* susceptibility *in vitro*, including for the slower-growing *M. restricta*.

## Results

First, we aimed to develop a novel experimental setup that allows for kinetic monitoring of *Malassezia* growth in a high-throughput manner, while limiting the need for specialized and costly lab equipment. We used flatbed scanners housed within a large incubator to automatically photograph *Malassezia* cultures in 96-well microtiter plates every two hours. With this setup, *Malassezia* MIC analyses were conducted using standard antimicrobial drug dilutions. By extracting the average brightness of each well, we were able to measure and quantify cell metabolism using resazurin dye as a growth indicator. This experimental protocol helped provide an unbiased and accurate relative MIC value for each drug condition tested. Upon optimization, we used this platform to study effects of itraconazole, isavuconazole, artemisinin and terbinafine on *Malassezia* species.

Overall, clinical isolates of *M. sloofiae* did not exhibit changes in susceptibility between strains. MIC50s of *M. sloofiae* were within one dilution step and exhibited the highest susceptibility to itraconazole (Table 1). *M. sympodialis* was highly susceptible to itraconazole with an MIC of 0.0098 μM and overall displayed drug susceptibility similar to *M. restricta* CBS7877, the only variation was a slight increase in susceptibility to isavuconazole, having a MIC50 value of 1.562μM compared to 0.0391μM (Table 1).

**Table 1:** MIC50 values for *Malassezia* strains. MIC50 is the first concentration for which growth has a relative growth of 50% or below. These values were calculated from the ratio of drug-treated cells to the growth of cells without drug treatment.

| Strain Number | Species | Itraconazole (µM) | Isavuconazole (µM) | Terbinafine (µM) | Artemisinin (µM) |
| --- | --- | --- | --- | --- | --- |
| CBS7877 | *M. restricta* | 0.0098 | 0.0391 | 0.3125 | 0.3125 |
| BTH004 | *M. restricta* | 0.0781 | 0.6250 | 0.3125 | 0.625 |
| KCTC27527 | *M. restricta* | 0.0098 | 0.0391 | 0.1562 | 0.625 |
| KCTC27524 | *M. restricta* | 0.0098 | 0.0781 | 0.1562 | 1.25 |
| BTH006 | *M. slooffiae* | 0.0391 | 0.1562 | 0.1562 | 0.625 |
| BTH007 | *M. slooffiae* | 0.0195 | 0.1562 | 0.1562 | 0.3125 |
| BTH016 | *M. slooffiae* | 0.0391 | 0.1562 | 0.1562 | 0.3125 |
| BTH012 | *M. sympodialis* | 0.0098 | 0.1562 | 0.3125 | 0.1562 |

For *M. restricta*, we examined the type strain CBS7877 and three *M. restricta* clinical isolates. Of the four antimicrobials, itraconazole was the most effective drug with a MIC range of 0.0098-0.0781 μM. The recent clinical strain BTH004 exhibited increased resistance to itraconazole relative to the wild-type strain, with a MIC50 of 0.0781 μM, 3 or 4 dilution steps higher.

Isavuconazole had a MIC range from 0.0781-0.625 μM. Isavuconazole resistance was observed in BTH004 which was four dilution steps more resistant than CBS7877 and KCTC27524 also exhibited resistance to isavuconazole. Terbinafine and artemisinin had higher MICs and were all within one dilution step: 0.6250 - 0.3125 μM. With the exception of KCTC27524 and KCTC27527 artemisinin MICs which both had a value of 2.5 μM.

**Figure 1:**
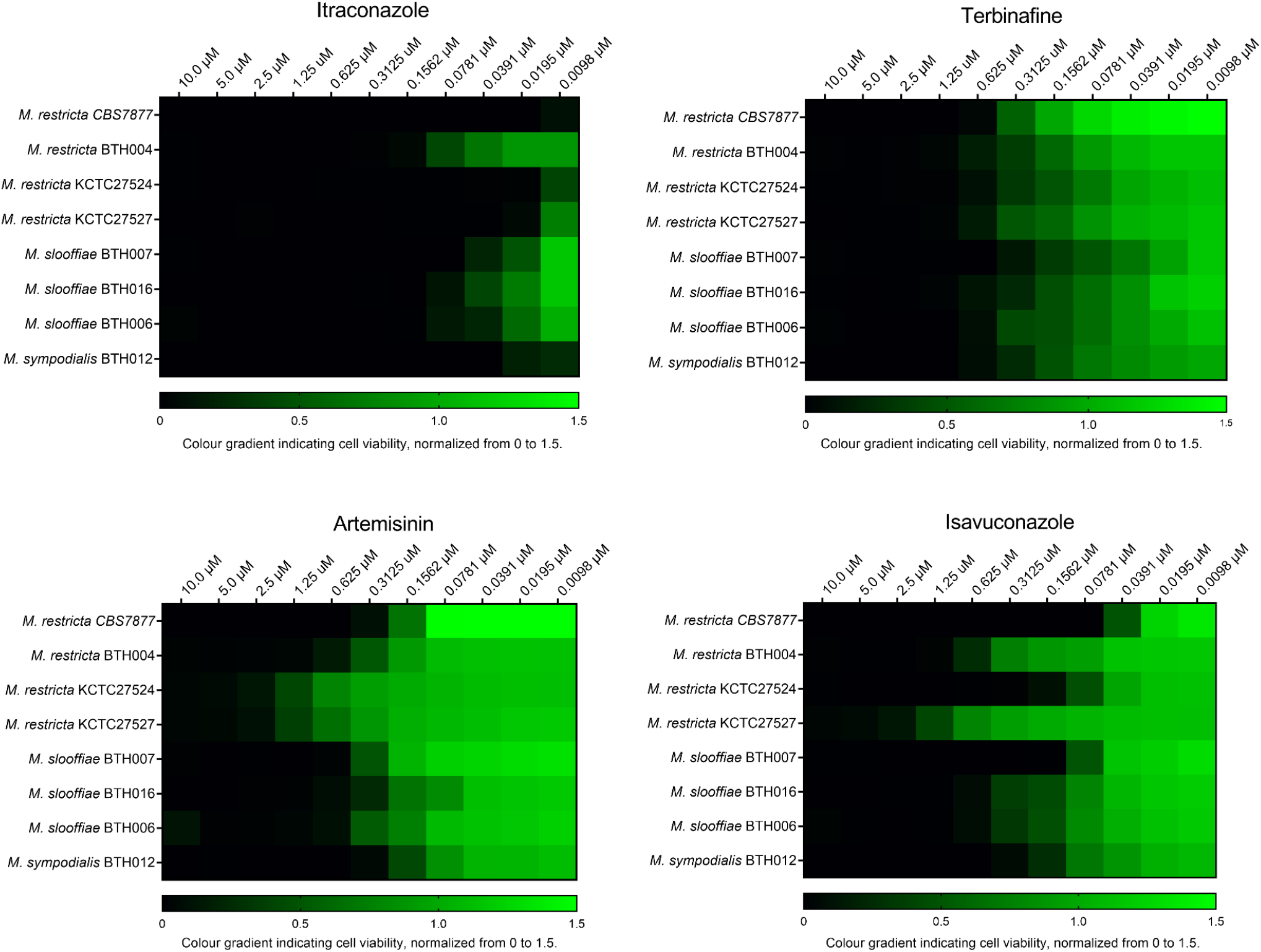
Heat map depiction of minimal inhibitory concentration (MIC) values from various *Malassezia* strains. Fungi viability values were created through computational quantification of luminescence, values were normalized from 0 to 1.5 with 1.5 being the average growth of the non-treated control wells.

## Conclusion

Here, we tested eight strains of *Malassezia* species against four antimicrobial agents and described the variable susceptibility to the antimicrobials tested. Overall, the antifungal itraconazole showed the highest potency against the diverse *Malassezia* strains studied. Resistance present between CBS7877 and at least one clinical *M. restricta* strain were present for isavuconazole, itraconazole and artemisinin, suggesting drug resistance development. Isavuconazole was FDA approved in 2015 while itraconazole has been used since 1992 and artemisinin has not been utilized against *Malassezia* infections. It is therefore unlikely that these strains have encountered either azole or artemisinin. Although artemisinin was less effective than the triazoles, it has been used in clinical applications with doses of up to 500 mg/day with no major side effects (20). Artiminisin resistance exhibited by both KCTC strains suggests resistance tied to pathogenic development as this drug has not been used to treat *Malassezia* ailments in the past. Therefore resistance is likely either tied to pathogenic development or developed from encountering other azole drugs utilized in treatment.

For the first time, we show both isavuconazole and artemisinin to be an effective agent against *Malassezia* species. These antimicrobials could be used as positive treatment options in the future. Our results agree with current literature that has suggested itraconazole to be an effective drug against the *Malassezia* genus (15, 16). However standardized testing for *Malassezia* species should be developed to enable the comparison of data between studies.

## Materials and Methods

### Fungal strains

*Malassezia* strains tested include *M. restricta (CBS7877), Malassezia* gifted from Won Hee Jung at the Chung-Ang University, Korea *M. restricta (KCTC27524, KCTC27527)* and *Malassezia* gifted from Bart Theelen from the Westerdijk Fungal Biodiversity Institute *M. restricta* (BTH 004), *M. slooffiae (BSS006, BSS007)*, and *M. sympodialis (BSS012, BSS016)*.

### Preparation of media

Leeming-Notman agar (LNA) Media was prepared by mixing 7.5g bacteriological grade peptone (Oxoid LP0037), 6g ox bile (Hardy Diagnostics C6511), 3.75g glucose, 0.075g yeast extract, 0.375g glycerol monostearate (Axenic 123-94-4), 0.375g chloramphenicol (Biobasic CB0118), and 3g of cycloheximide (Cayman Chemicals 14126), in 750ml of demineralized water, heated to 50-60°C then adding 375μl of glycerol, 750μl of Tween 60. Media was heated to 95°C before allowing it to cool, once cooled to 42C 0.03g of resazurin was added. Media was sterilized through a 0.045μm filter and stored at room temperature in a dark space for up to a week (stored for up to two weeks at 4°C).

### Preparation of fungal suspension

All *Malassezia* strains were cultured and maintained on LNA. Species identification was performed through Sanger sequencing internal transcribed spacer (ITS) regions then analyzed using NCBI-BLAST. Fungi were suspended in saline solution, to achieve a uniform inoculum, culture was suspended in bead bashing tubes and vortexed for 5 seconds. Inoculum was transferred to a glass tube, avoiding glass beads. Inoculum suspended in saline was then diluted to an OD600 of 0.3.

### Antimicrobial susceptibility testing

*Malassezia* activity was quantified through change of colour of the metabolic indicator dye resazurin. Antimicrobials tested (itraconazole, isavuconazole, artemisinin and terbinafine) all were dissolved in DMSO at a concentration of 2mM and stored at -20C. All protocols were performed in an Opentrons-OT2 liquid handler. In a 96-well flat bottom untreated NEST plate, each well was filled with 125ul of LNA-MIC media. Drugs were then added to column one to a concentration of 20uM and serially diluted across the plate until column 11. Fungal inoculum was added to 12mL of LNA-MIC and 155ul of inoculated media was added to each well. All wells are inoculated with fungi except for four negative control wells in column 12. Continuous monitoring allowed equal inoculation of each strain of fungi, regardless of species as end point times were readily altered. *Malassezia* strain was separately tested against all four drugs in duplicate in each 96-well microtiter plate.

Plates were covered with a gas permeable membrane (Diversified Biotech, BEM-1) and incubated at 34°C for 72 hours on a flatbed scanner (Canon CanoScan LiDE 300). Scans were performed automatically every two hours, and saved as 200 dpi color JPEG files. All wells start as dark blue, and slowly turn pink when resazurin is degraded, measuring the metabolic activity of *Malassezia* cells within the well. MIC results were recorded when the positive control wells fully flipped to pink (between 50 and 80 hours). Quantification of luminance was performed using a custom python script (tlscan2.py) which calculates the average brightness of each well using the Python Image Library (PIL). Briefly, each well is cropped out of the JPEG image using “.crop()”, then reduced to a single pixel using “.resize([1,1])” (this calculates the average brightness), then converted to grayscale using “.convert(‘L’)”, and finally extracted as a integer in the range 0 to 255 using “.getpixel((0,0))” (0 = darkest, 255 = brightest, encoded using sRGB gamma compression).

## Acknowledgments

We would like to thank members of the Shapiro lab for helpful discussions and technical assistance with this work. R.S.S is supported by a Tier II Canada Research Chair and a CIFAR Azrieli Global Scholars award. This work was funded by the Malassezia Foundation.

